# Integrating environmental DNA monitoring to inform eel (*Anguilla anguilla*) status in freshwaters at their easternmost range - A case study in Cyprus

**DOI:** 10.1101/2022.06.06.495005

**Authors:** Nathan P. Griffiths, Rosalind M. Wright, Bernd Hänfling, Jonathan D. Bolland, Katerina Drakou, Graham S. Sellers, Stamatis Zogaris, Iakovos Tziortzis, Gerald Dörflinger, Marlen I. Vasquez

## Abstract

**Aim:** Despite significant population declines and targeted EU regulations aimed at *A. anguilla* conservation, little attention has been given to their status at their easternmost range. This study applies wide scale integrated monitoring to uncover the present-day eel distribution in Cyprus’ inland freshwaters. These are subject to increasing pressures from water supply requirements and dam construction, as seen throughout the Mediterranean.

**Location:** Cyprus.

**Methods:** We applied environmental DNA metabarcoding of water samples to determine *A. anguilla* distribution in key freshwater catchments. In addition, we present this alongside ten years of electrofishing/netting data. Refuge traps were also deployed to establish the timing of glass eel recruitment. These outputs are used together, alongside knowledge of the overall fish community and barriers to connectivity, to provide eel conservation and policy insights.

**Results:** We confirm the presence of *A. anguilla* in Cyprus’ inland freshwaters, with recruitment occurring in March. Eel distribution is restricted to lower elevation areas, and is negatively associated with distance from coast and barriers to connectivity. Many barriers to connectivity are identified, though eels were detected in 2 reservoirs upstream of dams. The overall fish community varies between freshwater habitat types.

**Main Conclusions:** Eels are much more widespread in Cyprus than previously thought, yet mostly restricted to lowland intermittent systems. This makes for a case to reconsider the requirement for eel management plans. Environmental DNA based data collected in 2020 indicate that ‘present-day’ eel distribution is representative of 10-year survey trends. Suggesting that inland freshwaters may act as an unrealised refuge at *A. anguilla’s* easternmost range. Conservation efforts in Mediterranean freshwaters should focus on improving connectivity, therefore enabling eels to access inland perennial refugia. Thus, mitigating the impact of climate change and the growing number of fragmented artificially intermittent river systems.

## 1. Introduction

The European eel (*Anguilla anguilla*) is a catadromous fish species which spawns in the Sargasso Sea (Schmidt, 1923). Over the last four decades, significant global declines in eel populations have been observed (Aalto et al., 2016; Bilotta et al., 2011; Correia et al., 2018; Jacoby & Gollock, 2014; Podgorniak et al., 2016). *A. anguilla* recruitment is now estimated to be <10% of what was recorded in the 1970s (ICES, 2019; Trancart et al., 2020), and faces a myriad of pressures including migration barriers, habitat loss, turbine/pump mortality, overfishing/illegal exploitation, and climate change (Acou et al., 2008; Bilotta et al., 2011; Bolland et al., 2019; Buysse et al., 2015; Dekker, 2000; Moriarty & Dekker, 1997; Økland et al., 2019; Trancart et al., 2020). In response, the IUCN has classified *A. anguilla* as ‘critically endangered’ (Jacoby & Gollock, 2014), and the European Union has implemented specific legislation [The EC Eel Regulation (1100/2007)], requiring member states to develop eel management plans (EMPs). These regulations aim to facilitate the recovery of *A. anguilla* by implementing EMPs with the aim to achieve >40% of historic silver eel biomass (prior to anthropogenic impacts) safe passage (escapement) on their spawning migration between inland waters and the sea (Aalto et al., 2016; Council of the European Union, 2007). The European eel is distributed throughout the Mediterranean, and this region could provide a significant contribution to overall species recovery (Aalto et al., 2016). More recently, the General Fisheries Commission for the Mediterranean (GFCM) adopted the Recommendation GFCM/42/2018/1 (GFCM, 2018), establishing more targeted management measures for the European eel in the Mediterranean Sea (ICES, 2021). However, because the systematic investigation of eels in Cyprus’ freshwaters only commenced approximately a decade ago, present-day eel status on the island is subject to uncertainty (Zogaris et al., 2012).

Intermittent rivers and ephemeral streams (IRES), defined as watercourses which cease to flow at some point in time and space (Datry et al., 2018), are among the most common freshwater ecosystems globally (Larned et al., 2010). Furthermore, due to drying climates and increased human pressures on freshwaters, these systems are increasing in number, particularly in semi-arid regions (Datry et al., 2014, 2018). This has been noted in Mediterranean lotic systems, where increasing climate and anthropogenic pressures are leading to artificially intermittent river systems (Skoulikidis et al., 2011). The partial or complete drying of these systems is detrimental to whole fish communities, however upon flow resumption, fish recolonisation from upstream perennial reaches has been observed (Skoulikidis et al., 2011). The nature of these systems poses challenges for the implementation of eel management policy, since migration trigger flows may be present, but with no guarantee of perennial freshwater refuge (Zogaris et al., 2012).

Our case study area, the Mediterranean island of Cyprus, is part of the eastern limit of the European eels’ range. Until recently, quantitative eel catch data from the island’s freshwaters were lacking (Zogaris et al 2012). Therefore, the island was exempted from any obligation to implement eel management plans in 2009 (2009/310/EC). Through the Water Framework Directive (WFD) fish monitoring programme however, more recent data has suggested that *A. anguilla* may be more widespread in Cyprus than previously thought (Zogaris et al., 2012). Here, Zogaris et al (2012) found that eels were the most widespread native fish species remaining in the island’s freshwaters. Yet, when omitting data obtained from expert interviews and literature reviews, the observed *A. anguilla* occurrence was very low and localised. Cyprus represents a typical Mediterranean island, where lotic freshwater systems are often dominated by intermittent rivers and ephemeral streams (Papastergiadou et al., 2016). The historic natural state of most freshwater systems is not known, as anthropogenic impacts, namely water diversion and groundwater abstraction, have hugely impacted the freshwaters of the semi-arid island in recent years (Markogianni et al., 2014). Indeed, the inland freshwaters are influenced by an estimated 108 dams, one of the highest densities of dam reservoirs in Europe (Zogaris et al., 2012). Changes in flow regimes may force eels into summer refugia earlier, but with water retained upstream of likely impassible retention dams there is risk of restricting perennial habitat, while inducing artificial intermittence downstream (Skoulikidis et al., 2011). Despite these interruptions to natural flow regimes, there are catchments with perennial streams, particularly on the more humid western side of the island, and at higher elevations (Dörflinger, 2016; Papastergiadou et al., 2016; Zogaris et al., 2012).

While lentic inland freshwater habitat is present in Cyprus, natural wetlands are particularly vulnerable to climate change and anthropogenic impacts, yet artificial freshwaters are not generally managed with conservation values as a priority (Markogianni et al., 2014). A study by (Markogianni et al., 2014) found that most natural freshwater wetlands are smaller than artificial sites, and ephemeral in nature, with only 8 of the 30 largest wetlands classified as naturally occurring. Relating back to artificially intermittent streams (Skoulikidis et al., 2011), the presence of perennial wetlands have the potential to act as summer refuge habitat for fish species. This, of course, requires an element of - at least temporary - connectivity, and an assumption that wetland habitats are maintained year-round. Historically, such habitats were not valued as they were viewed as impediments to development and potential host-areas for disease (Markogianni et al., 2014). The current situation in Cyprus and other Mediterranean countries, is that natural wetlands are threatened by development and water use pressures (e.g. abstraction for irrigation), while artificial sites are not always managed with conservation values in mind.

In order to understand the value of Mediterranean freshwaters for eels, it is essential that we first have an improved understanding of the present-day eel status and distribution. To assess the eel distribution in Cyprus, and therefore inform current status, a wide range of catchments must be monitored in a short time frame. Environmental DNA (eDNA) metabarcoding of water samples has already proven to be effective for monitoring *A. anguilla* in heavily modified river systems in the UK (Griffiths et al., 2020). Therefore, this case study builds upon current knowledge by applying eDNA metabarcoding of water samples taken in 2020, alongside data from electrofishing/netting surveys carried out over a ten-year period (2009 - 2019) as part of the national WFD monitoring programme. By applying integrated monitoring methods, novel information on recent trends, and a snapshot of present-day fish distribution can be used to facilitate the improved understanding of *A. anguilla* distribution drivers. With multiple monitoring methods to inform this in such heavily managed and variable catchments, we aim to: a). Determine the current eel status and distribution in Cyprus; b). Assess the factors which influence eel distribution; and c). Consider the wider implications for Mediterranean freshwaters.

## 2. Methods

### 2.1 Environmental DNA metabarcoding (2020)

#### Sample collection

Twenty-six inland freshwater sites were targeted for eDNA surveys, with 5⨉ (1.5L) water samples taken from each between 04/02/2020 and 07/03/2020 (Figure 1). These sites were selected to include a representative range of freshwater habitat types, including; rivers (outlets) downstream of dams (n = 9), reservoirs upstream of dams (n = 9), unregulated rivers (n = 3), lentic wetlands (n = 3), and perennial spring-fed sites (n = 2) (Table S1). Based on recent assessments, and excluding reservoirs where water is retained by dams, only two wetland sites, both spring-fed sites, and one unregulated river were classified as perennial. The remaining systems have varying degrees of intermittence, although it is likely there are perennial refugia with unknown levels of connectivity in these catchments. Each of the five spatial replicate samples taken at each site consisted of 5 ⨉ 300ml sub-samples. A field/filtration blank was taken out into the field for each site, and processed alongside samples to monitor for contamination. Wherever possible, spatial replicates were taken at equidistant points spanning each study site, however where access did not allow this adhered to the following:

**Figure 1.**
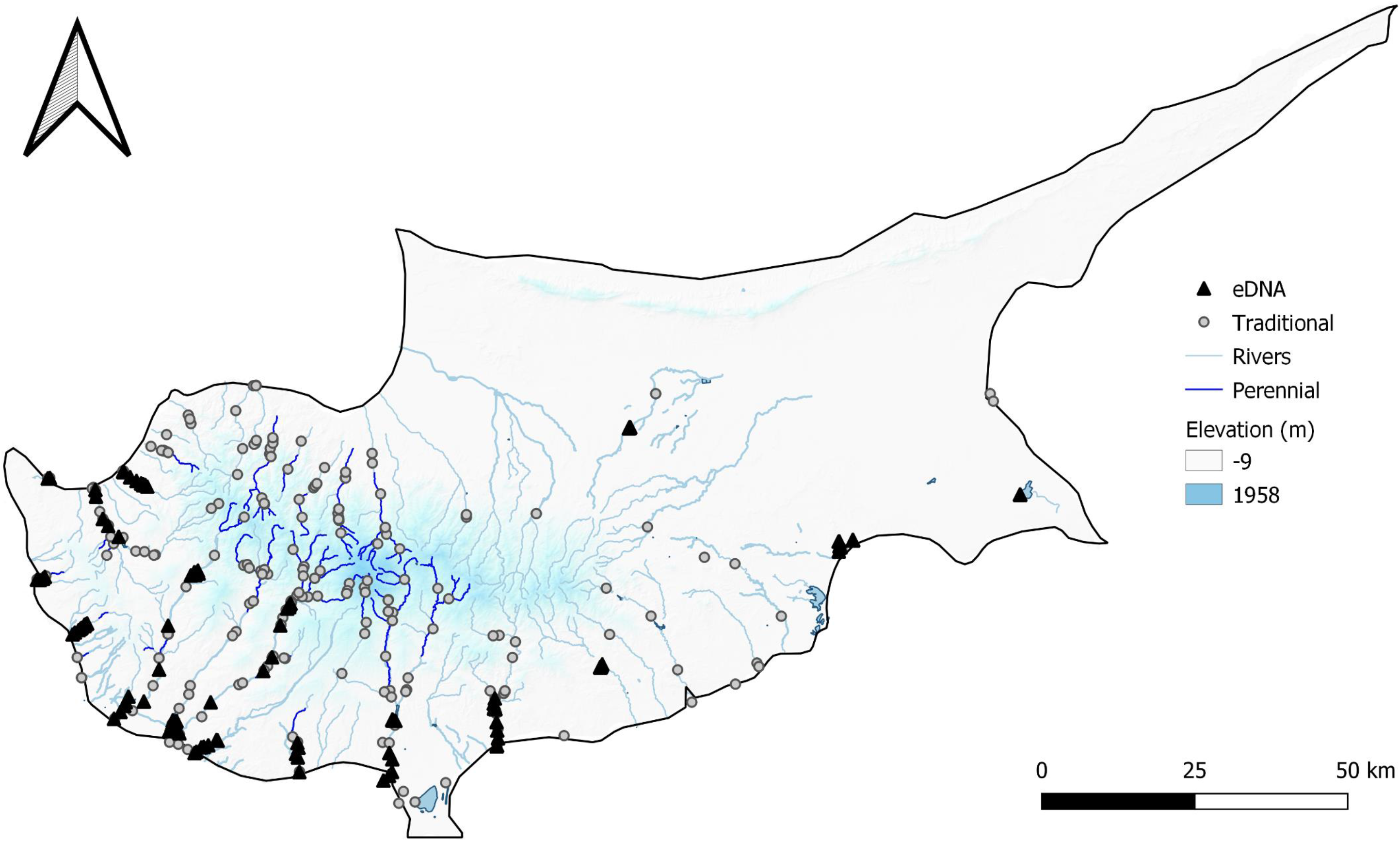
The distribution of data points across Cyprus, including the 130 eDNA sampling points in 2020 (black triangle) and the 299 fish survey points from 2009 - 2019 (grey dot). Areas of higher elevation are shaded in blue.

1. Rivers – Working from downstream to upstream, samples were taken at accessible points.

2. Reservoirs – When access was poor, sampling points were concentrated around the outlet where surface water was flowing suggesting increased mixing.

Measures were put into place in the field to avoid contamination, samplers wore sterile gloves while handling sampling equipment, which were changed between samples. Sampling bottles were only opened for the first time when at site prior to sampling, and samples were transported in a bleach sterilised coolbox on ice, to be filtered within 24 hours of collection. Following sample collection, water temperature was also recorded at each site (Table S1).

#### Filtration

Upon returning to the Cyprus University of Technology (CUT) laboratory, 1L of water from each sample was vacuum-filtered through sterile 0.45μm cellulose nitrate membrane filters with pads (47 mm diameter; Whatman, GE Healthcare, UK), using 2 filters per sample to reduce filter clogging. Prior to filtration all surfaces and equipment were sterilised with 10% bleach and 70% ethanol. Between filtration runs, filtration units were immersed in 10% bleach solution for 10 minutes, soaked in 5% v/v MicroSol detergent for an additional 10 minutes, and then rinsed thoroughly with purified water. Upon completion, filters were removed from units with sterile tweezers and placed back-to-back in 5ml polypropylene screw-cap tubes (Axygen, Fisher Scientific UK Ltd.), enclosed in sterile grip seal bags, and stored in a dedicated freezer at −20°C until DNA extraction. Filtration blanks (purified water) were processed alongside samples each day to monitor for contamination at this stage.

#### DNA extraction

DNA from duplicate filters was co-extracted following the Mu-DNA: Water protocol (Sellers et al., 2018), with minor changes resulting from the existing laboratory set up (Appendix I). As before, all surfaces were sterilised with 10% bleach and 70% ethanol, and all extraction reagents and plastics UV irradiated for 20 minutes prior to use. An extraction blank was included with each extraction run to monitor for contamination at this stage. To assess DNA quantity and purity, 2μl aliquots of each DNA sample were analysed on a Nanodrop 1000 Spectrophotometer (Thermo Fisher Scientific). Once completed, DNA extracts were stored at −20°C until PCR amplification.

#### Library preparation

Library preparation was carried out following an established 12S metabarcoding workflow previously developed at the University of Hull, the full protocol applied in this study can be viewed in Appendix I, however is summarised below:

Nested metabarcoding of DNA samples using a two-step PCR protocol was performed at CUT, using multiplex identification (MID) tags in the first and second PCR steps as described in (Kitson et al., 2019). PCR1 was performed in triplicate (3x PCR replicates per sample), amplifying a 106 bp fragment using published 12S ribosomal RNA (rRNA) primers 12S-V5-F (5′-ACTGGGATTAGATACCCC-3′) and 12S-V5-R (5′-TAGAACAGGCTCCTCTAG-3′) (Kelly et al., 2014; Riaz et al., 2011). Alongside DNA extracts PCR-negative controls (Molecular Grade Water) were used throughout, and positive controls (DNA (0.05 ng μl^-1^) from the non-native cichlid *Maylandia zebra*) were added to each plate outside of the eDNA prep area. All PCR replicates from each plate were pooled together to create sub-libraries and purified with MagBIND RxnPure Plus magnetic beads (Omega Bio-tek Inc., Norcross, GA, USA), following a double size selection protocol (Quail et al., 2009). Following this, a second shuttle PCR (PCR2) was performed on the PCR1 cleaned products to bind uniquely indexed Illumina adapters to each sub-library. A second purification was then carried out on the PCR2 products with the Mag-BIND RxnPure Plus magnetic beads (Omega Bio-tek Inc., Norcross, GA, USA). Eluted DNA was then stored at 4°C until quantification and normalisation. After normalisation and pooling, the final library was then purified again (following the same protocol as the second clean-up), quantified by qPCR using the NEBNext Library Quant Kit for Illumina (New England Biolabs Inc., Ipswich, MA, USA) and diluted to 4nM. The final library was then loaded (with 10% PhiX) and sequenced on an Illumina MiSeq using a MiSeq Reagent Kit v3 (600 cycle) (Illumina Inc., San Diego, CA, USA) at CUT.

Sub-library sequence reads were demultiplexed to sample level using a custom Python script. Tapirs, a reproducible workflow for the analysis of DNA metabarcoding data (https://github.com/EvoHull/Tapirs), was then used for taxonomic assignment of demultiplexed sequence reads. Reads were quality trimmed, merged, and clustered before taxonomic assignment against a curated national fish reference database. Taxonomic assignment used a lowest common ancestor approach based on basic local alignment search tool (BLAST) matches with minimum identity set at 98%. The full bioinformatics workflow is detailed in Appendix I.

### 2.2 Refuge Traps

Alongside eDNA sampling, refuge traps were placed near the tidal limits downstream of potential barriers to upstream migration in key freshwater catchments. These traps were made up from 2 x domestic mop heads (Pop Life, Paphos), tied together and secured in-stream. Refuge traps were checked regularly (every 2 to 4 days), lifted from the watercourse and emptied into a bucket to check for the presence of glass eels. These were deployed in key catchments and monitored from mid-March to mid-April 2019, and throughout January and February 2020.

### 2.3 National fish monitoring programme (2009 - 2019)

These data were primarily collected using in-stream backpack electrofishing surveys to determine the fish species present. In cases where electrofishing was not possible, nets were used to enable fish assessment. The final dataset over this period includes 299 spatial and temporal surveys, largely focussed in the more humid western region, but also covering more of the higher elevation central regions of the island (Figure 1). In addition to presence-absence data, the measurements of captured fish were taken, providing size data for the 355 individual eels. Since these surveys were carried out with WFD assessments in mind, they are not grouped into freshwater habitat type in the way eDNA surveys were, and so for the purpose of this study are considered as individual survey points.

### 2.4 Data analysis and visualisation

All downstream analyses were carried out using R version 3.6.3 (R Core Team, 2019). A low-frequency read threshold of 0.001 was applied to eDNA data, removing any reads which make up less than 0.1% of total reads assigned to any given sample as previously applied in other studies using this 12S marker (Griffiths et al., 2020; Handley et al., 2019; Hänfling et al., 2016). All maps were visualised, and associated metadata extracted using QGIS (QGIS Development Team, 2022). Data were screened for normality using base R. Since our data did not conform to a normal distribution, they were subsequently tested for associations using Spearman’s correlation tests (McDonald, 2014), and visualised using ggpubr (Kassambara, 2020). All other plots were visualised using ggplot2 (Wickham, 2016).

## 3. Results

### 3.1 eDNA metabarcoding

Following the application of quality controls, a total of 9,687,953 DNA reads were assigned to sample level, averaging 74,522 reads per sample. Of the 130 eDNA samples in this study, 31 (23.8%) were positive for *A. anguilla*, presenting relatively high eel reads (Range: 341 - 154303). The genus *Oreochromis* was omitted from our dataset due to identifying contamination in several blanks. Following this, across our 35 negative controls (negative, filtration blanks, and extraction blanks), 14 reads of *A. anguilla*, 14 reads of *Rutilus rutilus* and 4 reads of *Carassius* were detected in single filtration blanks. This is indicative of a low level of contamination, and we are therefore confident that our low-frequency reads threshold eliminates any risk of false positives arising in our dataset. This therefore confirms the presence of *A. anguilla* at 11/26 of our study sites (Table S1, Figure 2a). The 11 sites identified as positive for *A. anguilla* included four dam outlets, two reservoirs, two unregulated rivers, two spring-fed sites and one wetland (Figure 2a). The relative abundance and fish community detected is variable by site and freshwater habitat type (Figure 2). Combined site data show similarities in fish community composition between dam outlets and reservoirs (Figure 2b). Both are dominated by *R. rutilus* and share most other species, notably *A. anguilla* was detected with a higher relative read count and frequency below dams. The overall species community appears reduced by comparison in unregulated rivers, spring-fed, and wetland sites. Though it should be noted fewer of these habitat sites were sampled. Despite this, *A. anguilla* was detected in all freshwater habitat types (Figure 2a).

**Figure 2.**
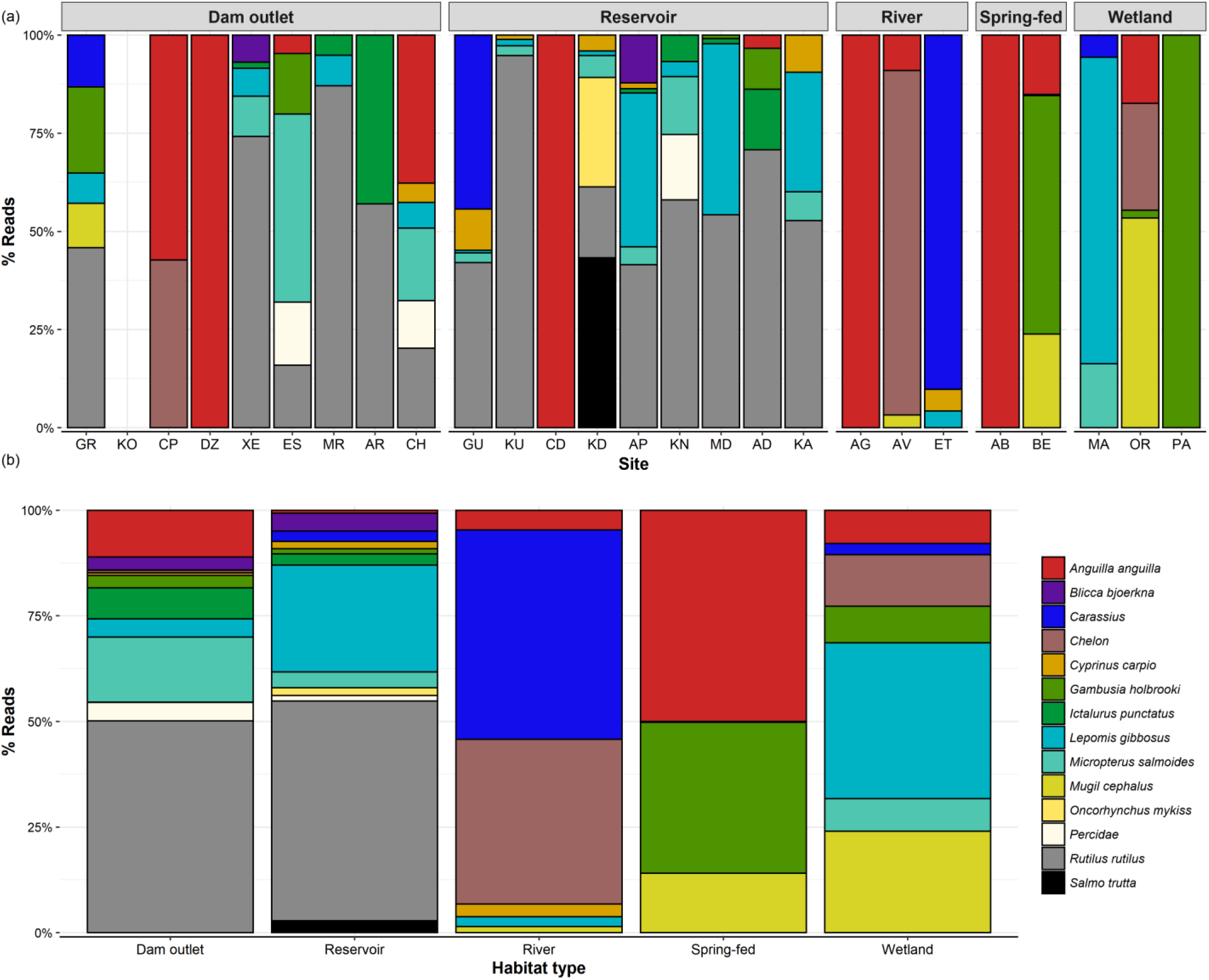
The relative abundance (% Reads) for each species detected, grouped into 5 freshwater habitat types. (a) The breakdown of eDN A survey sites within each habitat type. Note, that no fish DNA was detected at site KO. (b) Percentage reads merged by habitat type.

### 3.2 Fish monitoring programme

The 299 surveys captured a total of 355 *A. anguilla* specimens over the ten-year period (2009 - 2019). Of the eels captured during this period, a large proportion of individuals were caught in recent years (Figure 3). Though this may be, at least partly, due to progressive advances in knowledge of fish distribution and stream typology (Dörflinger 2016), thus leading to more targeted survey planning. Size classes suggest that the majority of eels captured were <50cm (Figure 3).

**Figure 3.**
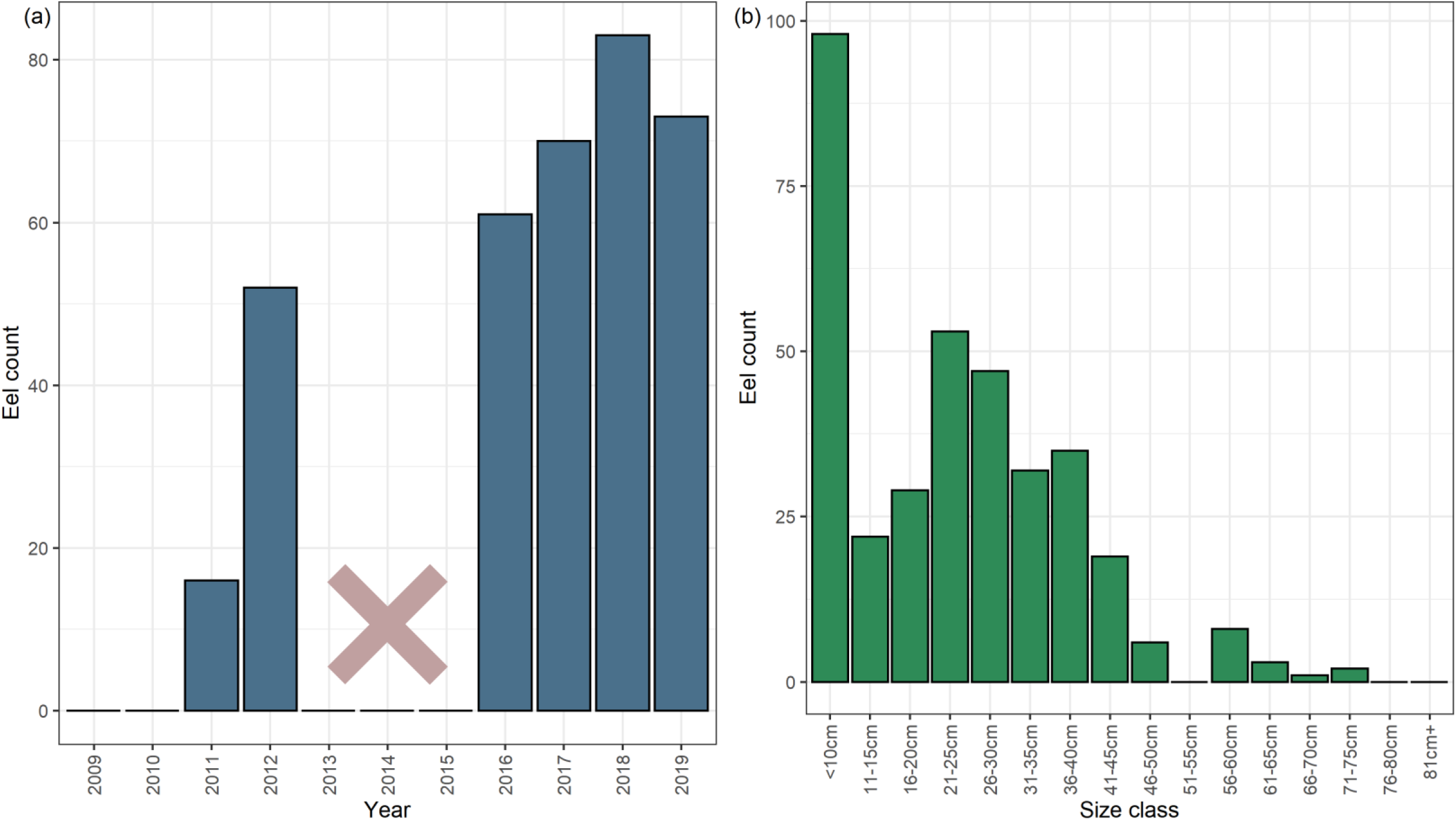
(a) The number of *A. anguilla* individuals captured each survey year, and (b) the size classes of *A. anguilla* captured. Note, that no fish surveys were carried out from 2013 - 2015, and no eels were captured in 2009 - 2010 surveys.

Overall, eel presence was confirmed in 61 of the 299 electrofishing/netting surveys (20.4%), mostly distributed in lowland systems on the western side of the island (Figure 4a). It should be noted however, that many of these positive surveys were due to repeat visits of the same catchments. Fish species richness appeared to be concentrated further inland, beyond the current distribution of *A. anguilla* (Figure 4b).

**Figure 4.**
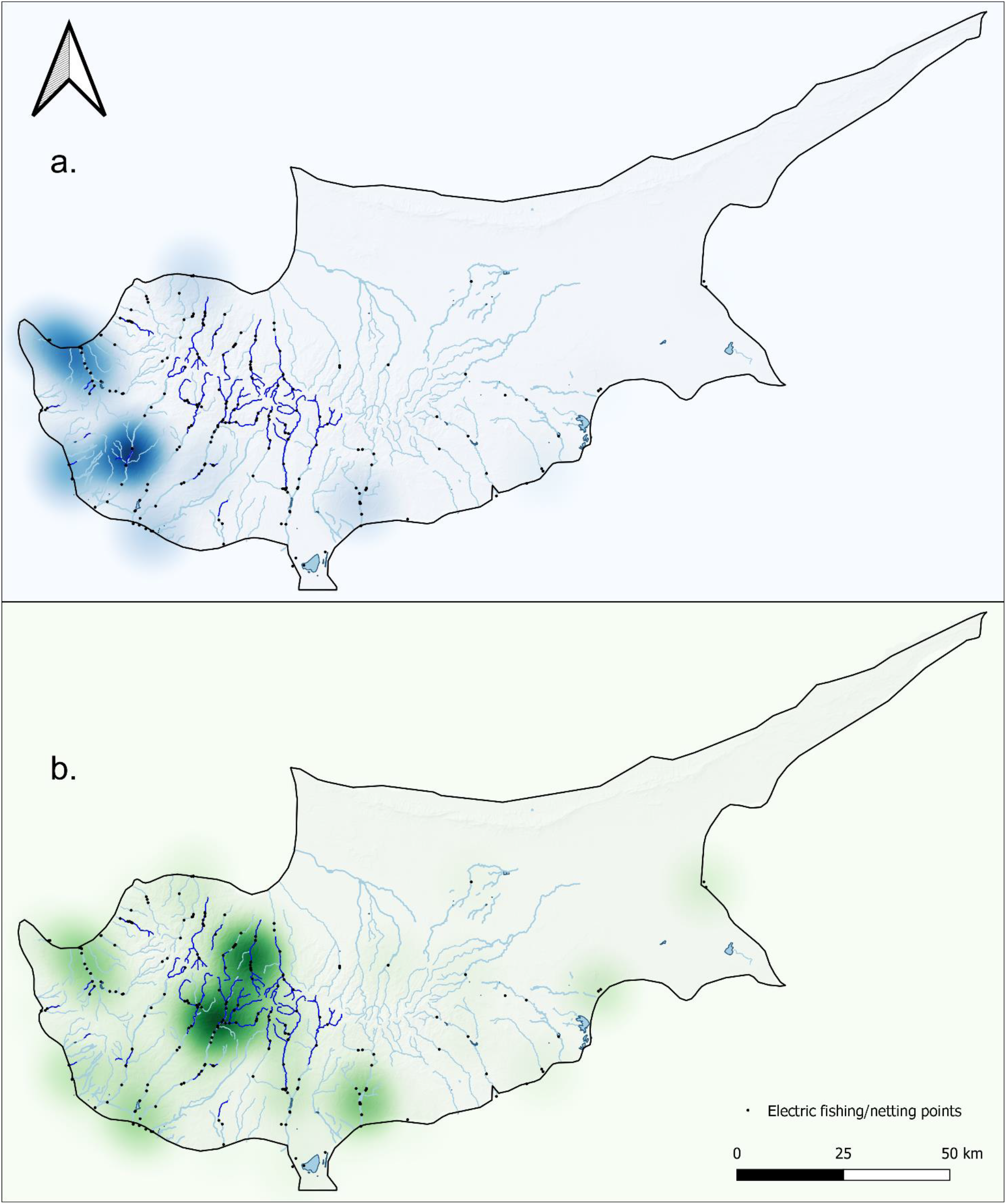
Heatmaps from fish surveys using Kernel Density Estimation, weighted by (a) eel catch and (b) species richness respectively.

### 3.3 Refuge traps

No glass eels were found in 2020, despite traps being regularly checked beyond our main eDNA sampling period (Table S1). However, in 2019 between 13/03/19 - 04/04/19, a total of 48 glass/pigmented eels were captured ranging from 50 - 66mm in length, confirming this method was effective when deployed in these catchments. This indicates that water sampling at key eDNA sites in 2020 occurred before eel recruitment that year.

### 3.4 Distribution of eels (combined methods)

Integrated monitoring gives us the best idea of overall eel distribution, and this shows a consistent pattern (Figure 5c). While both methods were able to detect *A. anguilla* up to 25km inland (Figure 5), water retention dams appear to be acting as barriers, preventing eels from accessing the central regions where overall species richness is highest (Figure 4). In these upper reaches, eel was present historically (Zogaris et al. 2012), but from the 299 fish surveys, only 1 eel was confirmed in this region in 2012 (Figure 5b). Excluding this record, and the 2/9 reservoirs positive for eel eDNA (Figure 2, Figure 5c), no other eels have been confirmed above dam structures. Both eDNA and catch methods found that eels were negatively correlated with elevation (R = - 0.33, p = <0.05; R = −0.4, p = <0.05), distance from coast (R = −0.35, p = <0.05; R = - 0.38, p = <0.05), and number of barriers (R = −0.39, p = <0.05; R = −0.45, p = <0.05) respectively (Figure 6). While overall species richness presented weak positive associations with the number of barriers, based on eDNA (R = 0.28, p = <0.05) and catch (R = 0.24, p = <0.05) methods. It is noted that the number of barriers, elevation, and distance from coast were all positively associated (Figure 6). Artificial instream barriers are widespread throughout Cyprus (Figure S1) (AMBER Consortium, 2020), yet their impacts have remained largely undocumented. In addition to the 108 dams, culverts (n=34) and weirs (n=42) are widespread on the island. While an additional 176 barriers are classified as ‘other’, suggesting no operational purpose, or that current information is lacking.

**Figure 5.**
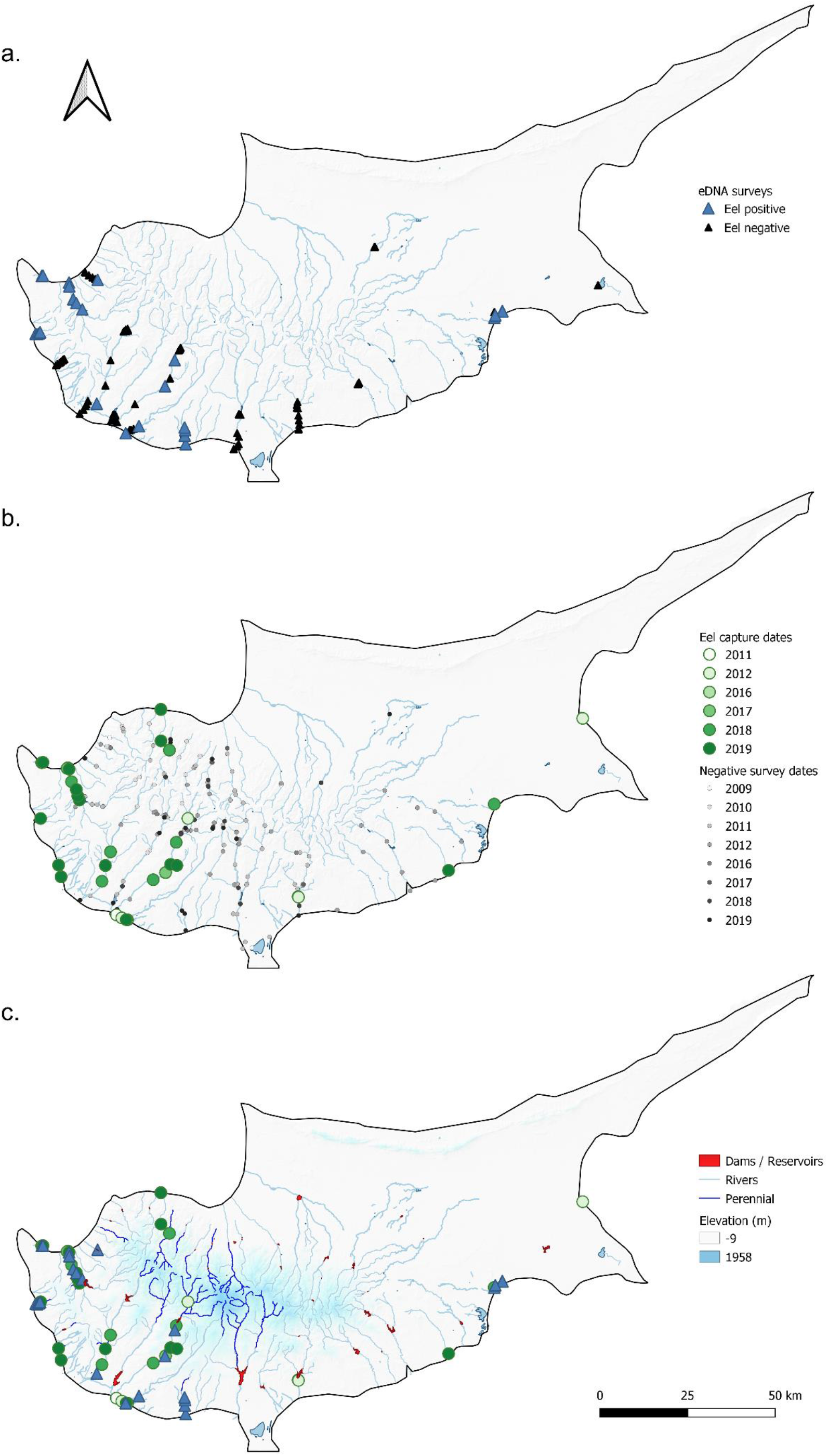
The distribution of *A. anguilla* based on (a) eDNA surveys, (b) electrofishing/netting, and (c) combined positive detections.

**Figure 6.**
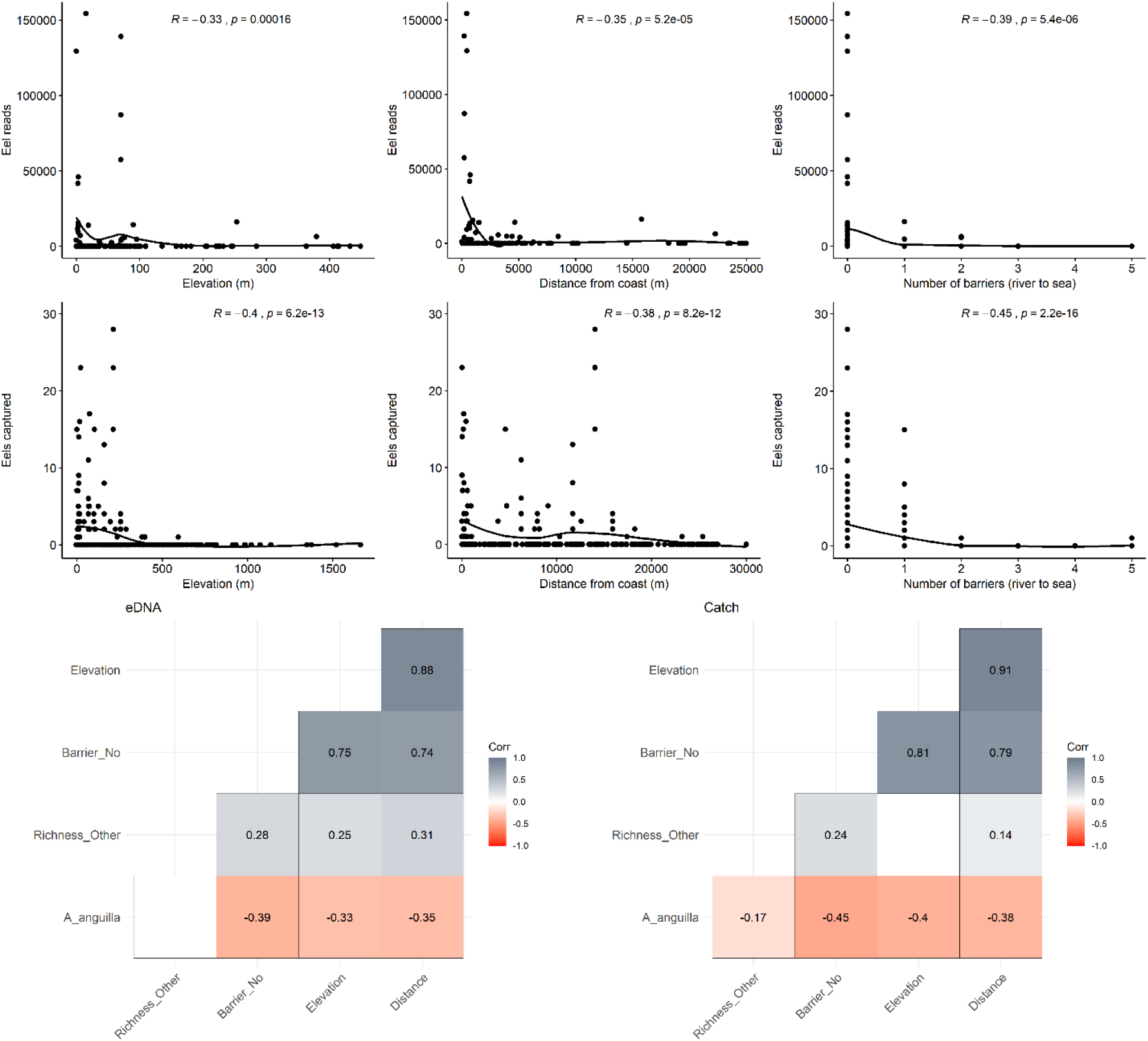
Scatter plots (top) including the Spearman’s rank correlation outputs, and smooth curves (loess) to visualise *A.anguilla* associations. Including a correlation matrix to visualise all eDNA based (bottom left) and catch based (bottom right) associations, blank values indicate no significance (p = >0.05).

## 4. Discussion

### 4.1. Eels in Cyprus

Our study confirms the presence of *A. anguilla* in Cyprus’ inland freshwaters. However, highlights that present-day eel distribution is restricted to lower elevation freshwater systems, primarily distributed in the western part of the island with relative close proximity to the coast (Figure 5). The freshwaters of Cyprus have largely been overlooked in regard to *A. anguilla* at policy level, and historically quantitative data regarding their distribution, presence and status have been lacking. We build on the initial study by Zogaris et al (2012), which reported eels were historically widespread in Cyprus based on literature reviews and expert interviews, but only physically captured/observed eels in three river basins. Our work applies integrated monitoring to expand on this, reporting present-day eel occupancy in a range of habitat types using eDNA monitoring (Figure 2), and widespread eel catches in the western lowlands (Figure 5). There was agreement between eDNA and catch surveys in terms of identifying eel ‘hot spots’, with two catchments in the west of the island consistently positive for *A. anguilla* (Figure 5c). With the knowledge that present-day eel distribution is restricted, we must consider that several streams historically documented as perennial have been recently observed to be in artificially intermittent states (Zogaris et al., 2012). This would suggest that, in recent years, the combination of barriers and widespread water retention upstream of dams is putting pressures on downstream lotic systems. Such pressures, including artificial intermittence, have led to fish kills in summer months. This was reported in 2013 (Zogaris, 2014) and 2021 (WDD, unpublished data), leading to high levels of eel mortality. This highlights that in the absence of access to perennial refugia, eels detected in lowland systems may be at risk. Similarly to our study (Figure 3), the majority of eels observed during these events were under 45cm in size (Zogaris, 2014). The data we present from electrofishing/netting surveys were collected after the spring influx of glass eels, and therefore may be biased towards same year recruitment. However, with so few eels captured measuring >50cm (Figure 3), there are indications that large female silver eels of the expected spawning size (∼65cm) remain rare (Clevestam et al., 2011). With that in mind, eDNA sampling was carried out before the elver run had peaked in the region (section 3.3). The eDNA detections from 2020 are therefore likely from eels which arrived in previous years, providing novel insight into the distribution of post-elver stage eels on the island.

The 2020 eDNA surveys detected eel presence in all habitat types (Figure 2). Including the detection of eels upstream of 2 reservoir dams (Table S1), this is novel information, since electrofishing is not carried out in reservoirs. Aside from these detections, and the one eel captured in 2012 in a stream above dam structures (Figure 5b), no other eels were found upstream of dams. Considered together, this provides evidence that while eels are entering Cyprus’ freshwater systems, they are likely restricted by dams and other barriers to lowland intermittent systems (Figure 6), where connectivity to summer refuge is limited and often restricted to relatively short spring-fed reaches (Dörflinger, 2016). These new insights highlight that despite historic records (Zogaris et al., 2012), eels are not currently present in the central and higher elevation regions of the island (Figure 5c, Figure 6), likely due to the many barriers to connectivity (Figure S1). While the central regions of the island were host to other fish species (Figure 4), it is apparent that currently, connectivity is not sufficient for eels to access them. Our eDNA and catch data both support this, since both methods found eel presence was negatively correlated to distance from sea, elevation, and number of artificial barriers (Figure 6). Eel presence is largely influenced by upstream passability (Teichert et al., 2020), and while eels are able to navigate some barriers to connectivity (Halvorsen et al., 2020), the cumulative impact of many barriers, in addition to large water retention dams, appears to be restricting their distribution. It is therefore not unexpected that more eels are found closer to the tidal limit, given their catadromous lifecycle.

Cyprus is a densely populated and semi-arid island, meaning that water stress on aquatic biodiversity is often unavoidable. As a result, the 108 water retention dams are relied upon to enable water management, with a capacity of 331,951,000m^3^ (WDD, 2017). With water scarcity considered an enduring issue, which may be further exacerbated by climate change, such pressures are hard to remove completely. There are, however, many artificial barriers to eel and fish passage, which may be more easily mitigated (AMBER Consortium, 2020) (Figure S1). By increasing connectivity within the lower reaches of catchments, the impact of upstream pressures may be diluted as more downstream habitat is available in the event water retention leads to an artificially intermittent environment in systems with previously perennial flows. Recent work by Dörflinger (2016) found the island’s river network is composed of 14% perennial and 86% intermittent stretches. Most perennial sites were classified as mountain streams, with an average elevation of 1051m, although fragmented stretches of perennial freshwaters at lower elevations are present. Until recently the distribution and extent of perennial streams was unknown, these improvements in knowledge of river typology will aid future planning of targeted monitoring and management actions.

Quantifying the passability and impacts of specific barriers was beyond the scope of this study. In the face of increasing pressure from barriers and anthropogenic impacts in Mediterranean freshwaters, up-to-date knowledge on site specifics is hard to feed into a robust management plan. Our data suggest that while barriers have a significant impact on eel distribution, in the absence of sufficient data, eel priority sites could be identified based on elevation and distance from coast (Figure 6). Based on our findings, we suggest that lowland intermittent systems with limited connectivity should be routinely monitored for eels as part of any future eel management plans. With the recommendation, that connectivity is increased between intermittent systems and perennial refuge, in addition to ensuring river to sea connectivity. This would increase the likelihood of eels accessing freshwater refuge, while allowing them to escape systems prone to desiccation and enabling silver eel migration.

### 4.2. Wider implications

Environmental DNA metabarcoding was able to detect our target species *A. anguilla* and fish communities in highly modified Mediterranean freshwater systems (Figure 2). In addition, the 2020 eDNA surveys were able to uncover widespread eel distribution, which broadly reflected the 10-year electrofishing surveys (Figure 5c). This highlights how eDNA metabarcoding can be applied successfully as part of an integrated monitoring programme in such systems. Allowing for a wide range of sites to be surveyed in a short time period, without compromising on detection rates. We suggest that eDNA metabarcoding of water samples would be a valuable tool in the implementation of any future Mediterranean eel management plans. By enabling wide scale sampling to identify eel status, and therefore focus limited resources on eel priority catchments. There are however some caveats to consider, mainly concerning the downstream transportation of eDNA. This can be observed in our eDNA data in this study, where due to downstream transportation of eDNA (Deiner et al., 2016; Milhau et al., 2019; Pont et al., 2018) and the potential for fish overspill/migration, reservoirs and their associated downstream outlets were found to have similar species compositions (Figure 2). When considering this for eel management, it is therefore important to interpret such data at a catchment scale. This explains why when plotting species richness spatially (Figure 4), only catch data are included, in order to ensure spatially precise distributions. There are also of course situations when physically handling the target species is required, in order to generate size class data for example. We therefore suggest that eDNA based monitoring is applied to systems with limited knowledge of eel distributions, with traditional surveys then targeting sites designated as eel priority. Combining both eDNA sampling and catch methods in an exploratory manner, to cover more terrain rapidly, may work best.

Our results are reflective of wider pressures on freshwater catchments in south and east Europe, where conservation action is required due to growing threats from river regulation, dam construction, hydropower and climate change (Szabolcs et al., 2022). Steps to identify eel priority sites, improve connectivity to water supply dams where possible, and consider eel passability in the design for future dams during the construction phase could be of benefit here. Given the recent GFCM recommendation GFCM/42/2018/1 (GFCM, 2018) to establish targeted management for the European eel in the Mediterranean Sea (ICES, 2021). Coupled with the knowledge that IRES are increasing in number in semi-arid regions (Datry et al., 2014, 2018), including the emergence of artificially intermittent river systems in the Mediterranean (Skoulikidis et al., 2011). The process of rethinking eel management plans in Cyprus, could be applied to inform other semi-arid regions in the development of future eel management strategies in freshwaters.

## 5. Conclusions

The findings of this study highlight the presence of *A. anguilla* in Cyprus’ freshwater systems, and identify that their present-day distribution is restricted. We present evidence that a critical problem is artificial river corridor fragmentation (Figure 6), caused by a high number of dams, weirs, culverts and other barriers (Figure S1). This therefore presents a case for the promotion of river connectivity restoration. Furthermore, the most stable and productive freshwater habitats are often upstream of dams (Figure 2, Figure 4), and therefore opportunities to improve eel passability here should be considered. Achieving this would in part take steps towards the implementation of an eel management plan, even if not in an official capacity. We therefore propose that targeted mitigation measures could be applied based on the data from this work. Prioritising fragmented lowland intermittent systems, with known eel presence. In addition, we propose environmental DNA metabarcoding of water samples as a method to complement wide scale fish monitoring programmes in the Mediterranean. Especially having in mind, the scattered and low-density eel populations in the regions’ freshwaters, eDNA methods can provide information on the presence of such populations in hard to sample areas. There has already been significant work put into the development of a fish bioindicator framework for Cyprus river basins (Zogaris et al., 2012). Effective monitoring of *A. anguilla* could be a key part of this, acting as an indicator of river-to-sea connectivity and well-connected perennial refuge areas further inland. This information is important at a broader scale, at the easternmost range of *A. anguilla*, where more localised eel management plans are now emerging (ICES, 2021). Environmental DNA based assessments of eel status could therefore help inform the status of inland freshwaters, while in addition allowing targeted eel management plans to be developed and implemented in areas sustaining present-day eel populations.

## Acknowledgements

We would like to thank the Water Development Department (WDD) in Cyprus for helping arrange access to sampling sites and providing data. We also give thanks to Robert Donnelly, Nektarios C. Efstathiou and Thessalia Nikolaou for laboratory support, and finally we thank Mick Bass for fieldwork support. Eel-targeted electrofishing and water sampling support was also provided by Dimitris Zogaris and Athina Papatheodoulou.

## Author Contributions

RMW, MIV, NPG, BH and JDB conceptualised this work. Writing was carried out by NPG, supported by RMW, MIV, BH and JDB. eDNA lab work and analysis was carried out by NPG supported by GSS, KD and MIV. Water collection was carried out by NPG, RMW, MIV, and electrofishing was led by SZ. WDD fish survey information and expert opinion of policy and local conditions was provided by IT, GD and SZ. All authors read, commented on, and approved this final manuscript.

## Data Availability Statement

Raw sequence reads have been archived on the NCBI Sequence Reads Archive (SRA) under BioProject ID: PRJNA844405.

All scripts and corresponding data have been archived and made available at Zenodo: https://doi.org/10.5281/zenodo.6617377.

