## Appendix I for "Integrating environmental DNA monitoring to inform eel (*Anguilla anguilla*) status in freshwaters at their easternmost range - A case study in Cyprus"

### Griffiths et al. Appendix I - Detailed eDNA methodology

#### eDNA capture and extraction

Water samples were vacuum-filtered within 24 hrs of collection in a dedicated eDNA area at Cyprus University of Technology (CUT). Surfaces and equipment were sterilised before, during, and after set-up in all work areas. Surfaces and vacuum pumps were wiped with 10% bleach solution. Non-electrical equipment was immersed in 10% bleach solution for 10 minutes, followed by 5% v/v MicroSol detergent (Anachem), and rinsed with purified water. Each 1 l water sample was filtered through sterile 0.45 µm mixed cellulose ester membrane filters with pads (47 mm diameter; Whatman, GE Healthcare) using manifold filtration units. Two filters were used for each sample and 30 mins allowed for water to pass through each filter, totalling a maximum of one hour filtration time per run. Blanks (n = 16) transported alongside samples were filtered during the last round of filtration for each sampling event. Equipment was sterilised after each round of filtration. Filters were removed from pads using sterile tweezers and placed in sterile 5 ml polypropylene screw-cap tubes (Axygen, Fisher Scientific UK Ltd.). Tubes were labelled, placed in grip seal bags, and stored at -20°C until DNA extraction. DNA extraction followed the Mu-DNA water protocol (Sellers et al., 2018). With the following adjustments due to existing lab set-up:

- Due to supply chain issues, guanidine thiocyanate was not included in our lysis buffer solution**

Briefly, captured DNA was liberated from the filter membranes via bead-milling lysis. The lysate underwent an inhibitor removal step prior to purification with silica membrane EZ-10 DNA Mini Spin Columns (NBS Biologicals) and final elution (100 µl). An extraction blank (n = 10), consisting only of extraction buffers, was extracted alongside samples for each run. 5 µl aliquots of each DNA extract were taken for measurement with a Nanodrop 1000 spectrophotometer (Thermo Fisher Scientific) to assess yield and purity. DNA extracts were then frozen at -20°C until PCR amplification.

#### eDNA metabarcoding library preparation

Dedicated rooms were available for pre-PCR and post-PCR processes. PCR reactions were set up in an ultraviolet (UV) and bleach sterilised laminar flow hood. Post-PCR processes were performed in a separate CUT laboratory. Libraries were prepared for sequencing using a nested metabarcoding workflow with a two step PCR protocol, where Multiplex Identification (MID) tags (unique 8-nucleotide sequences) were included in the first and second PCR for sample identification (Kitson et al., 2019). DNA extracts were PCR-amplified using vertebrate-specific primers that target a 106 bp fragment of the mitochondrial 12S ribosomal RNA (rRNA) region (Riaz et al., 2011). The primers were modified for the present study to include MID tags, heterogeneity spacers, sequencing primers, and pre-adapters. There were 24 unique MID tags for the forward and 24 unique MID tags for the reverse primers. This allowed each sample per library to be labelled with a unique forward and a unique reverse primer to reduce barcode misassignment and tag jumps (Deakin et al., 2014; Schnell et al., 2015). During the first PCR, samples were processed in sub-libraries, each including at least one filtration blank, one extraction blank, one negative control and one positive control. The PCR positive control was zebra mbuna (Maylandia zebra) DNA (0.05 ng/μl). M. zebra is not expected to be found in Cyprus' freshwater habitats.

The first PCR was performed in triplicate for each sample/control to combat stochasticity arising from low target DNA concentrations. PCR replicates for each sample/control had the same tag combination. Eight-strip PCR tubes with individually attached lids were used for PCR reactions. PCR reactions were performed in 25 μl volumes, consisting of: 12.5 μl of Q5® High-Fidelity 2x Master Mix (New England Biolabs), 0.5 μl of Thermo Scientific Bovine Serum Albumin (Fisher Scientific UK Ltd.), 7 μl of MGW (Fisher Scientific UK Ltd.), 1.5 μl of each 10 μM tagged primer, and 2 μl of template DNA. PCR reactions were sealed with mineral oil (Sigma-Aldrich) droplets. PCR was carried out with the following thermocycling profile: 98°C for 5 mins, 35 cycles of 98°C for 10 s, 58°C for 20 s and 72°C for 30 s, 72°C for 7 mins then held at 4°C.

PCR products were stored at 4°C until PCR technical replicates for each sample/control were pooled. 2 µl of each pooled PCR product was visualised on 2% agarose gels. PCR products were deemed positive where there was amplification at the expected size (200-300 bp) on the gel. PCR products were stored at 4°C until they were pooled according to band strength (no/very faint band = 20 µl, faint band = 15 µl, bright band = 10 µl, very bright band = 5 µl) on gel (Alberdi et al., 2018) to create sub-libraries for a double-size selection bead purification protocol. Ratios of 0.9x and 0.15x Mag-BIND RxnPure Plus magnetic beads (Omega Bio-tek) to 100 µl of each sub-library were used for purification. Eluted DNA (25 µl) was stored at 4°C until second PCR amplification.

The second PCR bound pre-adapters, MID tags, and Illumina adapters to the purified sub-libraries. Nine unique forward and reverse MID tag combinations were selected and applied to sub-libraries. Two replicates were performed for each sub-library in 50 µl reactions, consisting of: 25 µl of Q5 High-Fidelity 2x Master Mix (New England Biolabs), 13 µl of MGW (Fisher Scientific UK Ltd.), 3 µl of each 10 µM tagged primer (final concentration 0.6 µM; Integrated DNA Technologies), and 4 µl of template DNA. PCR was carried out with the following thermocycling profile: 95°C for 3 mins, 10 cycles of 98°C for 20 s and 72°C for 1 min, 72°C for 5 mins then held at 4°C. PCR duplicates for each sub-library had the same tag combination.

PCR products were stored at 4°C until duplicates for each sub-library were pooled. 2 µl of each pooled PCR product was visualised on 2% agarose gels. PCR products were deemed positive where there was amplification at the expected size (300-400 bp) on the gel. Sub-libraries were stored at 4°C until double-size selection bead purification. Ratios of 0.9x and 0.15x Mag-BIND RxnPure Plus magnetic beads (Omega Bio-tek) to 50 µl of each sub-library were used for purification. Eluted DNA (25 µl) was stored at 4°C until normalisation and final purification.

Sub-libraries were each quantified on a Qubit 3.0 fluorometer using a dsDNA HS Assay Kit (Invitrogen) and normalised by pooling according to sample size and library concentration. The pooled library was purified using the same ratios, volumes, and protocol as the second PCR purification. Based on the Qubit™ concentration, the library was diluted to 4 nM. The library was then quantified by qPCR using the NEBNext Library Quant Kit for Illumina (New England Biolabs). Based on the qPCR concentration, the library was adjusted to 4 nM and denatured following the Illumina MiSeq library denaturation and dilution guide. The final library was sequenced at 13 pM with 10% PhiX Control on an Illumina MiSeq using 2 x 300 bp V3 chemistry (Illumina).

#### Bioinformatics (Tapirs + database curation)

#### Reference database

To represent commonly detected European fish species, the curated UK fish species reference database of (Hänfling et al., 2016) was used as the basis for the creation of the Cyprus reference database used in this study. A species list of all recorded fish caught in Cyprus inland freshwaters was used to determine fish species required in the original database. Records that included the 12S mitochondrial region for these listed species were downloaded from GenBank and used to update the original curated database. Duplicate sequences and poor quality references (those with ≥ 100 ambiguous bases) were removed prior to final BLAST database creation.

#### Cyprus species list:

*Anguilla anguilla*
*Aphanius fasciatus* *
*Atherina boyeri* *
*Blicca bjoerkna*
*Carassius auratus*
*Carassius carassius*
*Chelon saliens* *
*Cyprinus carpio*
*Dicentrarchus labrax* *
*Gambusia holbrooki* *
*Gobius paganellus* *
*Hypophthalmichthys molitrix*
*Ictalurus punctatus* *
*Lepomis gibbosus*
*Leuciscus aspius* (!)
*Micropterus salmoides*
*Mugil cephalus* *
*Nerophis ophidion* *
*Oncorhynchus mykiss*
*Oreochromis aureus* *
*Oreochromis niloticus* *
*Perca fluviatilis*
*Rutilus rutilus*
*Salmo trutta*
*Syngnathus abaster* *
*Chelon labrosus* *
*Chelon ramada* *

“*” Cyprus specific database, (!) no GenBank record

#### Taxonomic assignment

Sequencing data was automatically demultiplexed to separate (forward and reverse) fastq files per library using the onboard Illumina MiSeq Reporter software. Library sequence reads were further demultiplexed to sample using a custom Python script. Tapirs, a reproducible workflow for the analysis of DNA metabarcoding data (https://github.com/EvoHull/Tapirs), was used for taxonomic assignment of demultiplexed sequencing reads. Tapirs uses the Snakemake workflow manager (Köster & Rahmann, 2012) and a conda virtual environment to ensure software compatibility.

Raw reads were quality trimmed from the tail with a 5 bp sliding window (qualifying phred score of Q30 and an average window phred score of Q30) using fastp (Chen et al., 2018), allowing no more than 40% of the final trimmed read bases to be below Q30. Primers were removed by trimming the first 18 bp of forward and reverse reads. Reads were then tail cropped to a maximum length of 106 bp and reads shorter than 90 bp were discarded.

Sequence read pairs were merged into single reads using fastp, provided there was a minimum overlap of 20 bp, no more than 5% mismatches and no more than 5 mismatched bases between pairs. Only forward reads were kept from read pairs that failed to be merged. A final length filter removed any reads longer than 110 bp to ensure sequence lengths approximated the expected fragment size (~106 bp).

Redundant sequences were removed by clustering at 100% read identity and length (--derep_fulllength) in VSEARCH (Rognes et al., 2016). Clusters represented by less than three sequences were omitted from further processing. Reads were further clustered (--cluster_unoise) to remove redundancies due to sequencing errors (retaining all cluster sizes). Retained sequences were screened for chimeric sequences with VSEARCH (--uchime3_denovo).

The final clustered, non-redundant query sequences were then compared against the reference database using BLAST (Zhang et al., 2000). Taxonomic identity was assigned using a custom majority lowest common ancestor (MLCA) approach based on the top 2% query BLAST hit bit-scores, with at least 90% query coverage and a minimum BLAST hit identity of 98%. Of these filtered hits, 80% of unique taxonomic lineages therein had to agree at descending taxonomic rank (domain, phylum, class, order, family, genus, species) for it to be assigned a taxonomic identity. If a query had a single BLAST hit it was assigned directly to this taxon only if it met all previous MLCA criteria. Read counts assigned to each taxonomic identity were calculated from query cluster sizes. Lowest taxonomic rank was to species and assignments higher than order were classed as unassigned.
