## Supplementary material for "Integrating environmental DNA monitoring to inform eel (*Anguilla anguilla*) status in freshwaters at their easternmost range - A case study in Cyprus": Figure S1

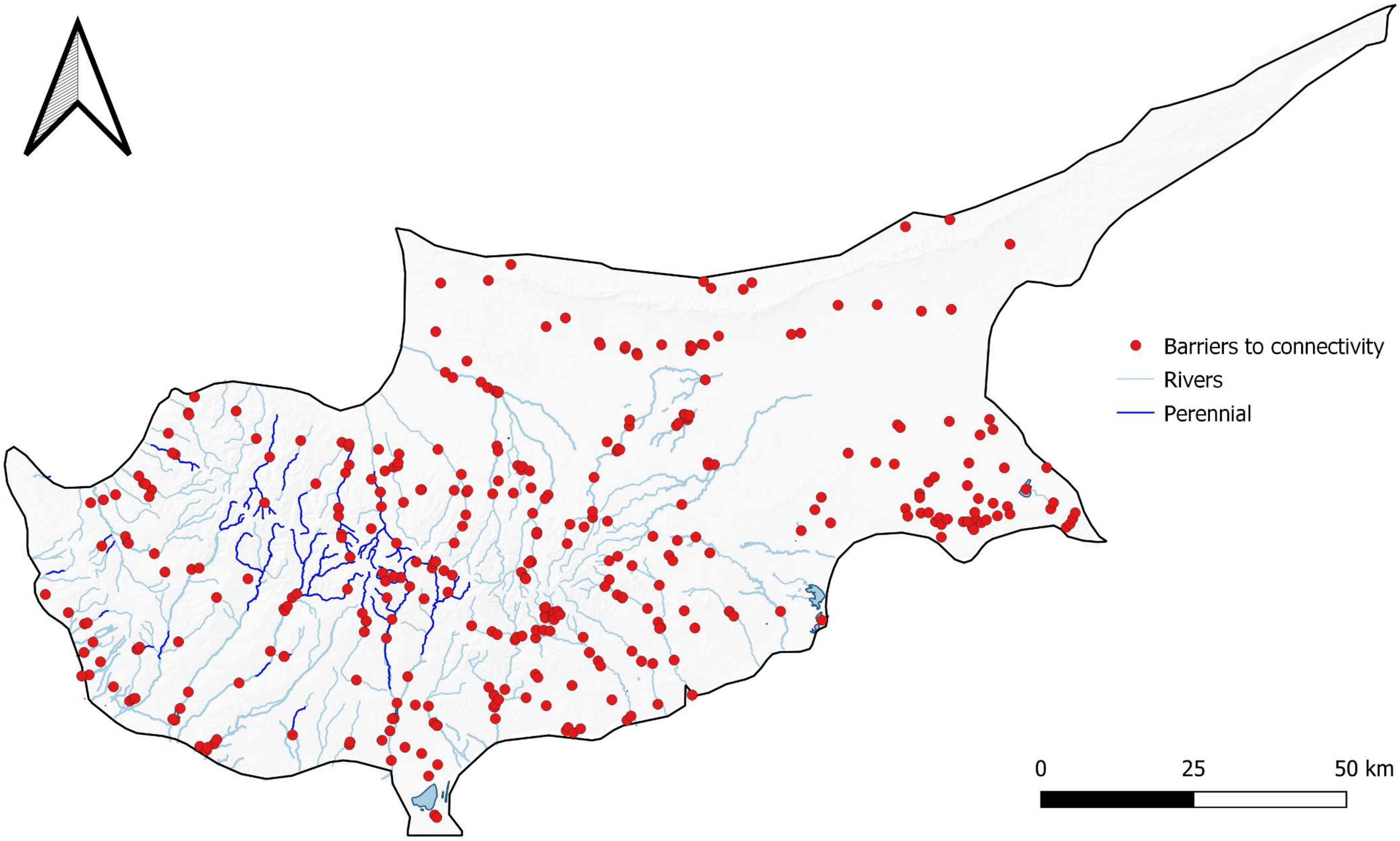


**Figure S1.** The distribution of artificial instream barriers in Cyprus, based on data from the AMBER Consortium (2020).
